# Disruption of the NTCP-EGFR receptor complex as a strategy for selective inhibition of hepatitis B and D virus entry

**DOI:** 10.64898/2026.02.05.703960

**Authors:** Atsuto Kusunoki, Kaho Shionoya, Frank Stappenbeck, Takeshi Morita, Hirofumi Ohashi, Makoto Nagano, Ryo Morishita, Feng Wang, Kazuhiko Katayama, Farhad Parhami, Koichi Watashi

## Abstract

Hepatitis B and D virus (HBV, HDV) enter hepatocytes through a coordinated process mediated by a receptor complex consisting of sodium taurocholate co transporting polypeptide (NTCP) and its entry cofactors, including epidermal growth factor receptor (EGFR). Here, we established an in vitro assay to evaluate the NTCP-EGFR interaction and identified Oxy229, an oxysterol-based compound that disrupted this molecular interaction. Oxy229 selectively inhibited HBV and HDV infection to HepG2-NTCP cells and primary human hepatocytes. Mechanistic analysis revealed that Oxy229 impaired the relocalization of the HBV-NTCP complex from plasma membrane to intracellular vesicles. Notably, Oxy229 did not compromise the physiological functions of NTCP and EGFR, i.e., bile acid transport and activation of downstream EGFR signaling pathways including Ras-MAPK and PI3K-Akt pathways, indicating selective inhibition of viral entry. Compound derivative analysis identified Oxy283, which acquired dual inhibitory activity against both NTCP-EGFR interaction and NTCP multimerization, resulting in enhanced anti-HBV potency. These findings establish the functional significance of the NTCP-receptor complex formation in HBV/HDV entry and highlight this machinery as a potential target for antiviral intervention.

## Introduction

Hepatitis B virus (HBV) infection constitutes a major global health burden, an estimated 250 million people chronically infected worldwide (1). Chronic HBV infection elevates the risk of developing liver diseases, including liver cirrhosis and hepatocellular carcinoma (2). Hepatitis D virus (HDV) is a satellite virus coinfected with HBV in an estimated 5% of chronically HBV-infected people and is associated with the most severe clinical symptom of viral hepatitis (2). For developing new anti-HBV and HDV agents, it is vital to understand the molecular mechanisms underlying virus infection.

HBV carries three envelope proteins, large (LHBs), middle (MHBs), and small (SHBs) surface antigens in virions. HBV entry into hepatocytes follows a multiple-step process: HBV attaches to host cells initially in a non-specific manner through heparan sulfate proteoglycans on cell surface, and exploits a more specific and high affinity interaction of LHBs, via its N-terminal segment called the preS1 region, especially its 2-48 aa region, with sodium taurocholate cotransporting polypeptide (NTCP), a hepatocyte-specific bile acid transporter that serves as the host entry receptor (3, 4). HDV caries the HBV-derived surface antigens on its virions and thus is regarded to follow the same or similar NTCP-dependent entry mechanism as HBV. This virus-NTCP interaction is therapeutically targeted by bulevirtide (also known as Myrcludex B), a preS1-mimicking peptide that competes for NTCP binding to inhibit HBV/HDV infection and is clinical approved for treatment to HBV/HDV-coinfected patients (5). Successful development of bulevirtide has demonstrated the therapeutic significance of the viral entry process as a drug target. However, efforts for discovery of anti-HBV/HDV entry inhibitors have largely focused on targeting the viral surface antigens and NTCP itself, leaving other host factors involved in the entry underexplored.

Although NTCP is essential for hepatocyte-specific HBV/HDV attachment, NTCP alone is insufficient to mediate viral entry. We have previously shown that epidermal growth factor receptor (EGFR) functions as an entry cofactor by interacting with NTCP and supporting the receptor function (6, 7). Upon stimulation with ligands such as EGF, EGFR is phosphorylated at multiple tyrosines and recruits the adaptor molecules that trigged EGFR endocytosis, leading to the internalization of the HBV/HDV-bound NTCP-EGFR complex (defined as viral receptor complex) from cell surface to intracellular vesicles. Consistent with this role, an EGFR tyrosine kinase inhibitor, Gefitinib, was shown to suppress HBV and HDV infection by blocking EGFR phosphorylation and subsequent internalization of the viral receptor complex, suggesting EGFR as a potential antiviral target. However, Gefitinib also blocks the original function of EGFR that activates the downstream signaling such as the Ras-MAPK and PI3K-Akt pathways, raising concerns about potential adverse effects. Thus, alternative strategies that disrupt the entry cofactor function but minimize the impact on the essential EGFR signaling would be desired. We have also reported that NTCP in the multimerization form facilitated HBV internalization through yet unknown mechanism (8, 9). Troglitazone and its derivative were identified to inhibit NTCP multimerization and HBV internalization, although their activities were relatively modest (9, 10). So far, approach of the viral receptor complex as an anti-HBV/HDV target has not been well documented.

In this study, we primarily focus on the interaction between NTCP and EGFR, the core of the viral receptor complex, as a target of compound discovery. By developing an in vitro NTCP-EGFR binding assay and using this system for chemical screening, we identified Oxy229, a semi-synthetic oxysterol that interrupted the NTCP-EGFR interaction and found this compound to selectively inhibiting the infection of HBV and HDV. Oxy229 inhibited the EGFR-mediated internalization of the preS1-NTCP complex, without affecting the EGFR downstream signaling and NTCP-dependent bile acid transport. We further generated a derivative, Oxy283, which interestingly acquired the dual inhibitory activity to both NTCP-EGFR interaction and NTCP multimerization, thereby showing enhanced antiviral potential. These findings show the significance of the viral receptor complex for HBV/HDV entry, and as a potential target for anti-HBV/HDV agents.

## Results

### Oxy229 inhibits the binding between NTCP and EGFR

To evaluate the molecular interaction between NTCP and EGFR, we established the AlphaScreen assay that can quantify the specific binding between NTCP and EGFR in vitro. Using the same methodological strategy that we reported previously to evaluate the HBV LHBs-NTCP interaction (11), we synthesized recombinant His-tagged NTCP (NTCP-His) and biotinylated EGFR (EGFR-bio) proteins in the wheat germ cell free protein expression system, and incubated together, followed by adding streptavidin-conjugated donor beads and anti-His antibody-conjugated acceptor beads (Fig. 1A, upper): In this assay, proximation of the two proteins induces the transfer of singlet oxygen from the donor beads to the acceptor beads and produces AlphaScreen signal. Actually, co-incubation of NTCP-His and EGFR-bio produced AlphaScreen signal 21-times higher than co-incubation of a non-relevant His-tagged glutathione S-transferase (GST), instead of NTCP-His, and 24 times higher than co-incubation of a control biotinylated dihydrofolate reductase (DHFR), in place of EGFR-bio (Fig. 1A, lower), indicating the specific binding between NTCP and EGFR. Using this assay, we screened 1,000 compounds by quantifying the AlphaScreen signal upon treatment with each compound at 100 μM (see Materials and Methods in detai1) (Fig. 1B, upper) and identified 40 compounds that reduced the signals to less than 50% of the DMSO-treated controls. Then to eliminate false positive compounds, including those inhibiting either 1) production of AlphaScreen signal itself, 2) transfer of singlet oxygen, and 3) binding of biotin to streptavidin, or of His to anti-His antibody, we performed the secondary AlphaScreen assay using biotinylated-His protein, which constitutively produced AlphaScreen signal independently of protein-protein binding (Fig. 1C, upper). This signal is inhibited by false positive compounds but unaffected by NTCP-EGFR interaction inhibitors. By selecting compounds that reduced the signal in the primary screening (Fig. 1B) but not the secondary screening (Fig. 1C), we identified Oxy229 (Fig. 2A) as one of the hit compounds showing the highest effect to reduce the NTCP-EGFR interaction-dependent signal with relevant data reproducibility. Oxy229 exhibited a dose-dependent reduction in the AlphaScreen signal produced by coincubation of NTCP-His and EGFR-bio (Fig. 1B, lower), whereas this compound did not significantly affect the signal produced by bio-His protein (Fig. 1C, lower). These data suggest that Oxy229 specifically inhibits the interaction between NTCP and EGFR.

**Fig. 1.**
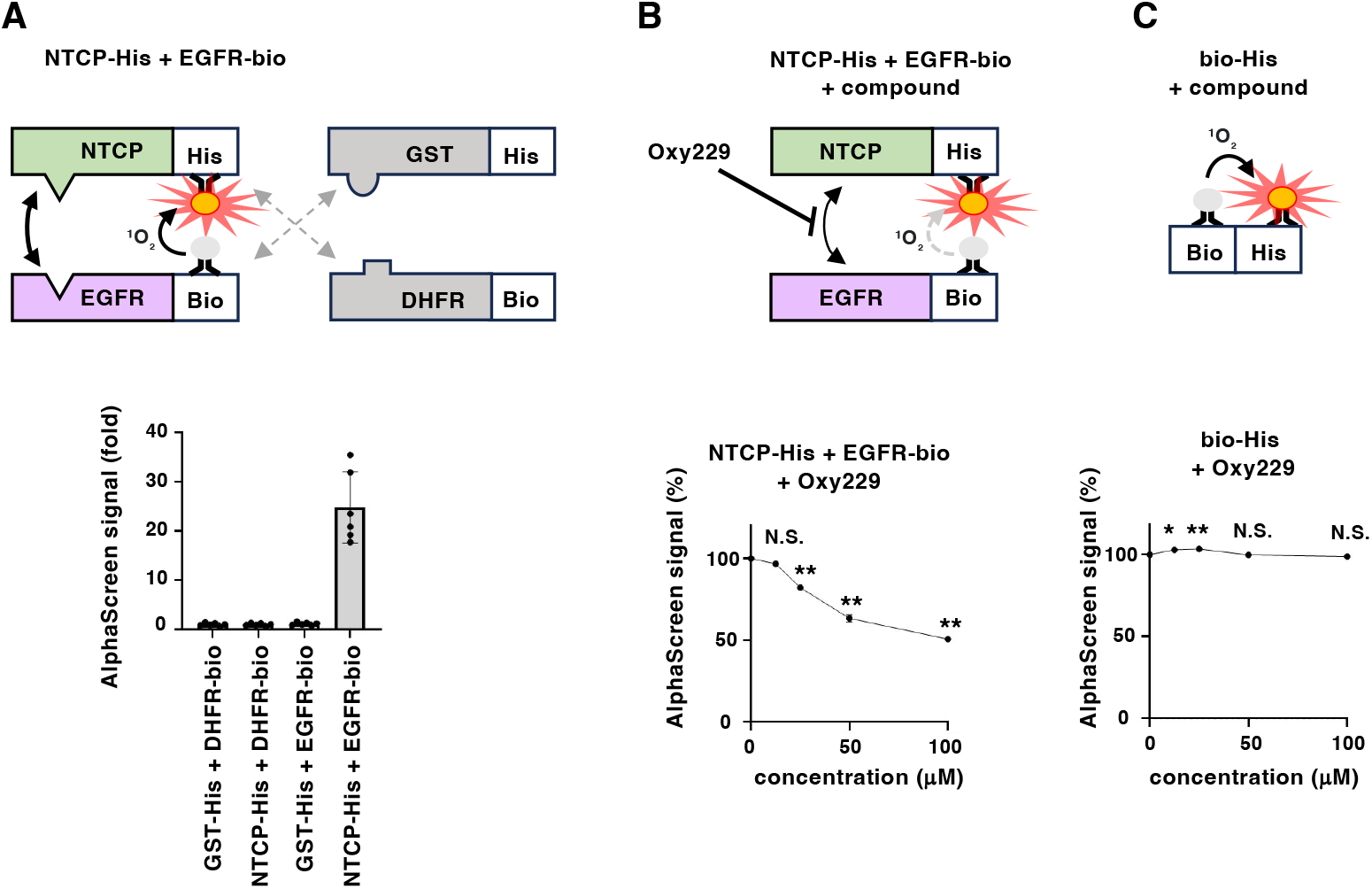
Oxy229 inhibited the NTCP-EGFR interaction in vitro. **(A)** Upper, schematic diagram of the AlphaScreen assay evaluating the interaction between His-tagged protein and biotinylated protein in vitro. Recombinant His-tagged NTCP (NTCP-His) or GST (GST-His) as a non-relevant control and biotinylated EGFR (EGFR-bio) or DHFR (DHFR-bio) as a control are incubated with streptavidin-coated donor beads and anti-His-conjugated acceptor beads. Proximation of biotinylated protein and His-tagged protein induces transfer of singlet oxygen from the donor beads to the acceptor beads and produces the AlphaScreen signal. Lower, AlphaScreen signal by incubation with NTCP-His (or GST-His as a contro1) and EGFR-bio (or DHFR-bio as a contro1) are shown by graph. **(B)** Upper, schematic diagram of the assay evaluating the effect of the compounds on NTCP-EGFR interaction in vitro. Lower, AlphaScreen signals in the presence or absence of Oxy229 (12.5, 25, 50, and 100 μM), are shown. **(C)** Upper, schematic diagram of the secondary assay to neglect false positive compounds, in which the AlphaScreen signals constitutively produced by biotinylated-His tag peptide (bio-His) are detected. Lower, AlphaScreen signals in the presence or absence of Oxy229 (12.5, 25, 50, and 100 μM) are shown. Statistical significance was determined by One-way ANOVA (**, P<0.01; *, P<0.05; N.S., not significant).

**Fig. 2.**
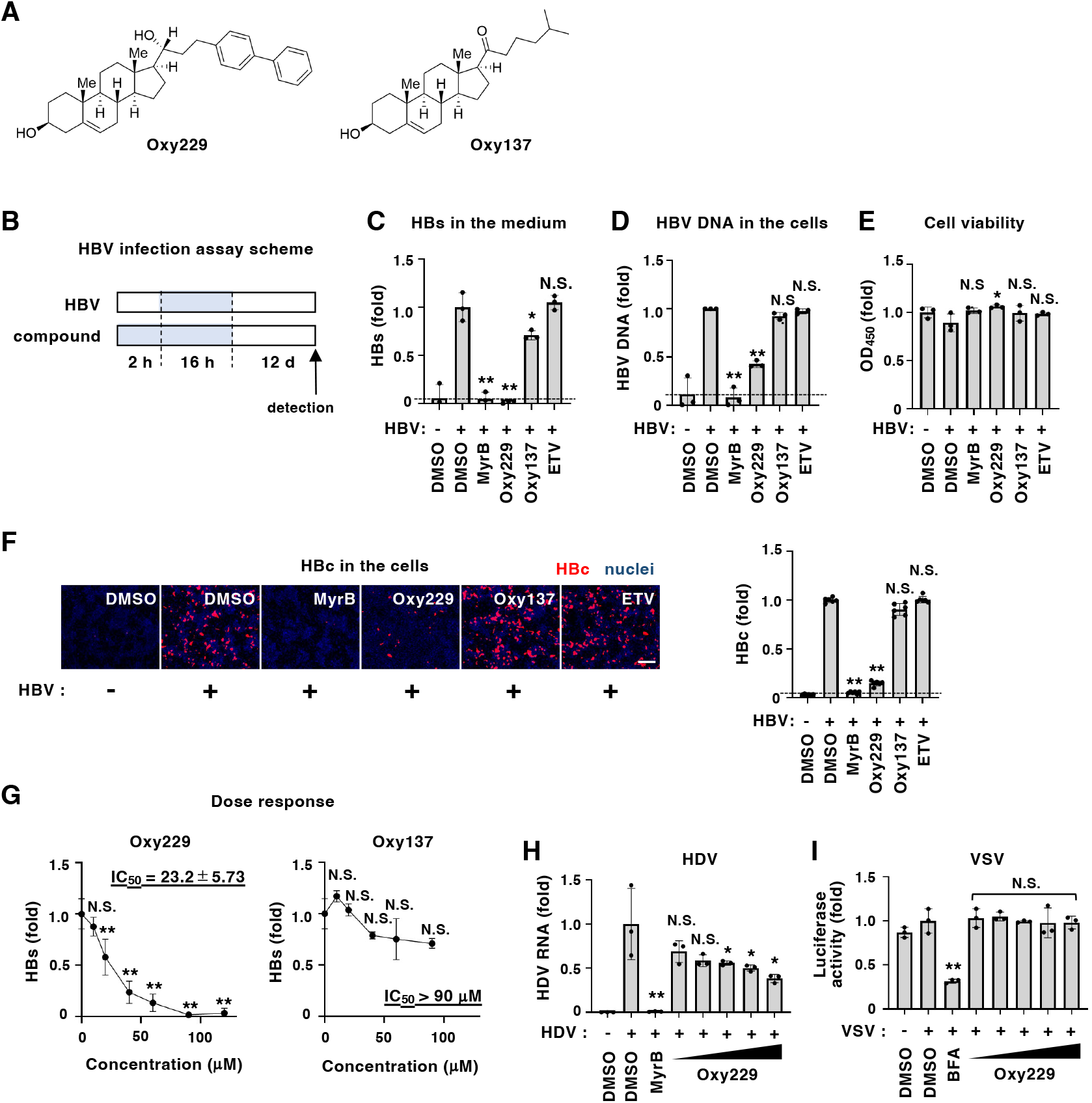
Oxy229 inhibited HBV and HDV infection. **(A)** Chemical structures of Oxy229 (left) and its derivative, Oxy137 (right). **(B)** Schematic representation of the schedule for HBV infection assay. HepG2 cells overexpressing NTCP (HepG2-hNTCP-C4 cells) were treated with compounds for 2 h and inoculated with HBV along with the compounds for 16 h. After washed out free HBV and compounds, the cells were cultured in the absence of compounds for 12 days to detect HBV infection markers, HBs antigen in the cultured supernatant and HBV DNA and HBc antigen in the cells. Cell viability was also examined to evaluate the cytotoxicity of compounds. Blue and white bars indicate the presence or absence of HBV or compounds. **(C-F)** HBV infection assay was performed as shown in (B) in the presence or absence of the indicated compounds [0.9% DMSO, 100 nM Myrcludex B (MyrB), 90 μM Oxy229, 90 μM Oxy137, 500 nM Entecavir (ETV)]. HBV infection was determined by detecting HBs antigen in the cultured supernatant (C), HBV DNAs (D) and HBc antigen (F) in the cells. Cell viability was measured by WST assay (E). Fluorescence areas for HBc antigen (red signa1) were quantified and are shown as a graph on the right in (F). Scale bar, 100 μm. **(G)** HBV infection assay was performed with various concentrations of Oxy229 (left) and Oxy137 (right) (0, 10, 20, 40, 60, 90 and 120 μM). 50% maximal inhibitory concentrations (IC_50_s) are also shown. **(H)** For evaluation of HDV infection, HepG2-hNTCP-C4 cells pretreated with the compounds for 2 h were inoculated with HDV along with the compounds for 16 h. After washing and culturing cells for five days, intracellular HDV RNA was measured by real time RT-PCR. **(I)** VSV-mediated entry was evaluated by inoculating Huh-7 cells with VSV pseudovirus for 1 h, followed by washing and incubating cells for 72 h to measure the luciferase activity. Statistical significance was determined by One-way ANOVA (**, P<0.01; *, P<0.05; N.S., not significant).

### Oxy229 specifically inhibits HBV and HDV infection

We then evaluated the effect of Oxy229 on HBV infection in a cell-based HBV infection assay: HepG2-NTCP-C4 cells were pretreated with the compounds for 2 h and were inoculated with HBV in the presence of the compounds for 16 h, followed by washing out free virus and incubating the cells for 12 days to detect HBs antigen in the culture supernatant by ELISA, HBc antigen in the cells by immunofluorescence, and HBV DNA in the cells by real time PCR as HBV infection markers, as well as cell viability by WST assay (Fig. 2B). We used Myrcludex B (MyrB) an anti-HBV/HDV entry inhibitor (12), as a positive control and entecavir (ETV), a clinically-approved HBV replication inhibitor (13) as a negative control. In this infection assay, MyrB drastically reduced HBs, HBc, and HBV DNA, but ETV showed no significant effect on these infection markers (Fig. 2C-F), indicating that this assay can evaluate the effect of HBV entry inhibitor, consistent with the published reports (14, 15). Similarly, Oxy229 significantly reduced HBs, HBc, and HBV DNA, without showing cytotoxicity (Fig. 2C-F). This anti-HBV activity of Oxy229 was dose-dependent, showing HBs reduction up to 10% of the DMSO-treated control, and its 50% inhibitory concentration (IC_50_) was calculated to be 23.2 ±5.7 μM (Fig. 2G, left). By contrast, Oxy137, a control compound, sharing the same cholesterol backbone as Oxy229 but a different sidechain (Fig. 2A, right) having no significant effect on NTCP-EGFR interaction (see Fig. 5C), and did not show apparent reduction in HBV infection (Fig. 2C-G), which was used as an inactive (or less active) reference compound in this study.

We further confirmed that Oxy229 inhibited HBV infection to primary human hepatocytes, as shown by reduction in HBs and HBc, without cytotoxicity, as the case with MyrB and an EGFR antagonist, Gefitinib (7), used as another positive control (Fig. S1). Oxy229 also inhibited the infection of HDV in a dose-dependent manner (Fig. 2H), which follows NTCP-and EGFR-dependent cell entry (6, 7), whereas it did not significantly affect the entry of vesicular stomatitis virus (VSV), as an NTCP-independent virus (Fig. 2I). Thus, Oxy229 specifically inhibited HBV and HDV infection.

### Oxy229 inhibits the internalization of HBV from the plasma membrane

We then examined which step in the HBV lifecycle was inhibited by Oxy229 (Fig. S2A). The HBV lifecycle is divided into 1) the entry process, including the viral attachment to the target cells, internalization inside the cells, and membrane fusion to nuclear translocation for the formation of covalently closed circular DNA (cccDNA), a template of virus replication, followed by 2) the replication process, including transcription to viral RNAs, encapsidation into nucleocapsids, reverse transcription to generate genomic relaxed circular DNA (rcDNA), and viral release outside of the cells (16) (Fig. S2A). We at first evaluated the HBV replication activity using Hep38.7-Tet cells, which is a Tet-Off cell system including HBV replication from a chromosome-integrated HBV transgene by depletion of tetracycline (17). The cells were treated with the compound upon depletion of tetracycline continuously for six days to detect HBV DNA produced in the culture supernatant by real-time PCR. Whereas HBV DNA levels were drastically reduced by ETV, used as a positive control, but not MyrB or Gefitinib, as negative controls, HBV DNA was not significantly affected by Oxy229 or Oxy137 (Fig. S2B). We then evaluated preS1-mediated viral attachment by treating HepG2-hNTCP-C4 cells with a TAMRA fluorescence-labeled preS1 peptide consisting of the receptor binding region spanning 2 to 48 aa in the HBV preS1 region to detect cell surface-bound TAMRA fluorescence (preS1-TAMRA), as previously reported (7) (Fig. S2C, upper). In this preS1 attachment assay, treatment with MyrB used as a positive control for an attachment inhibitor, but not Gefitinib as a negative control internalization inhibitor (7, 12), completely reduced the preS1-mediated cell attachment (Fig. S2C-c and f). In contrast, Oxy229 and Oxy137 did not significantly affect preS1-cell attachment (Fig. S2C-d and e). The following HBV internalization activity was examined using the same fluorescence peptide: HepG2-hNTCP-C4 cells were treated with preS1-TAMRA at 4ºC for 30 min for allowing attachment, followed by transfer the cells to 37ºC upon EGF stimulation to initiate internalization in the presence or absence of the compounds for 60 min, to observe the subcellular localization of TAMRA fluorescence with a confocal microscopy (Fig. 3A, upper). Whereas preS1-TAMRA was detected mainly on cell surface without 37ºC incubation (Fig. 3A-a), preS1-TAMRA was relocated with speckled patterns inside the cells, as well as the cell surface localization, after 37ºC incubation upon DMSO treatment as a control (Fig. 3A-b). Remarkably, these internalized preS1 speckles were rarely observed but stayed on the cell surface by treatment with Oxy229, but not Oxy137 (Fig. 3A-d and e). Cell number quantification indicated that the cell population exhibiting internalized preS1 speckles was clearly decreased from approximately 70% in the DMSO treated control to 5% in Oxy229 treated cells. (Fig. 3A, upper right graph). These data suggest that Oxy229 inhibits the HBV internalization.

**Fig. 3.**
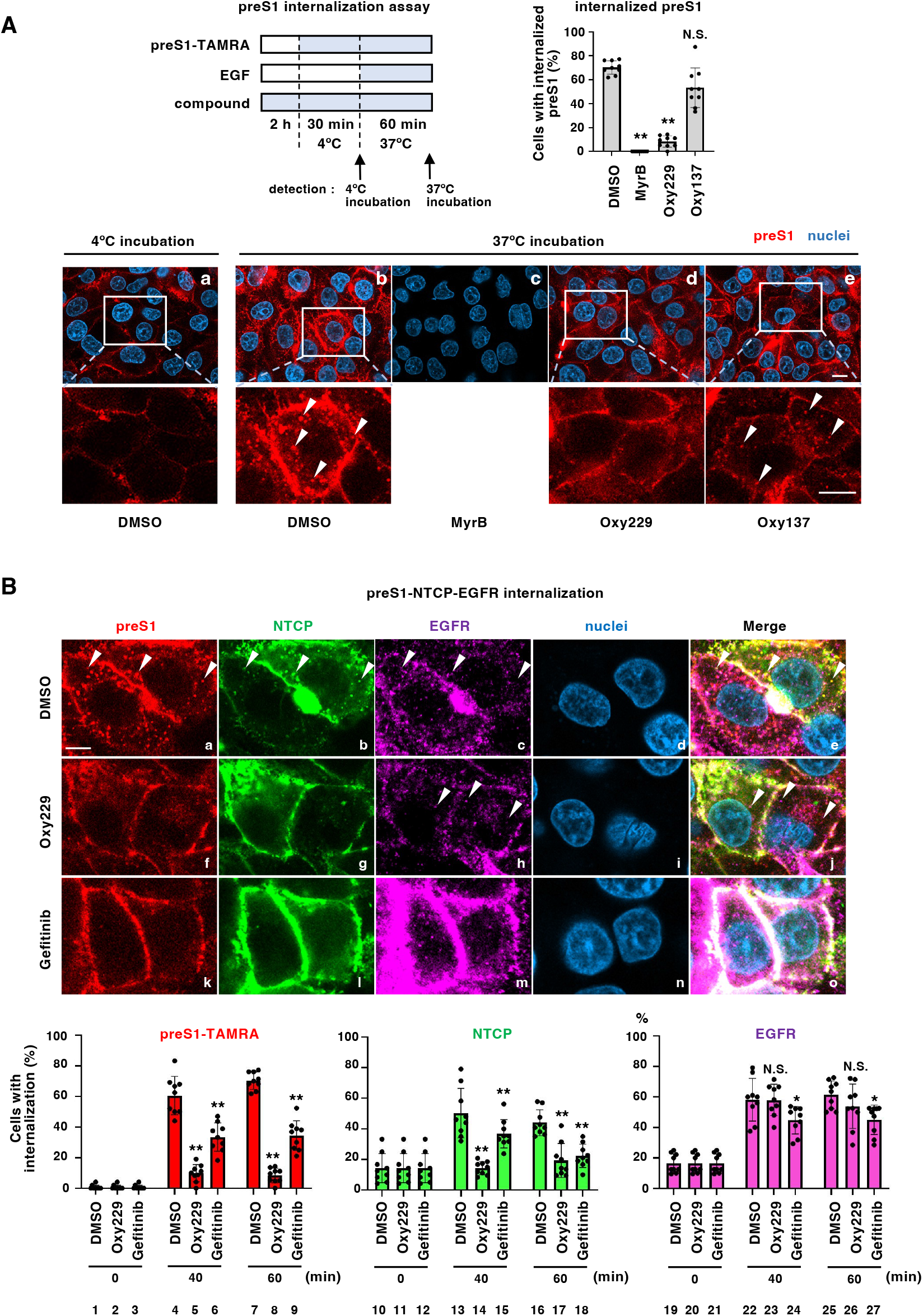
Oxy229 inhibited HBV internalization. **(A)** HBV internalization was evaluated with TAMRA-labeled N-terminally myristoylated preS1 peptide consisting of 2-48 amino acid region (preS1-TAMRA). HepG2-hNTCP-C4 cells treated with preS1-TAMRA at 4ºC for 30 min (to allow preS1-cell attachment) were incubated upon 10 ng/ml EGF at 37ºC for 60 min (to induce preS1 internalization) in the presence or absence of the indicated compounds. preS1-TAMRA (red) as well as nuclei (blue) were observed with a confocal microscopy. Insets in the upper pictures (white boxes) are indicated at the lower pictures with higher magnification. White arrows at the lower pictures show the internalized preS1 speckles. Internalization of preS1 was quantified and is shown as a graph (upper right). Scale bar, 10 μm. **(B)** Subcellular localizations of preS1 (red), NTCP (green), and EGFR (purple) were examined in HepG2-hNTCP-oxSG cells that were treated as shown in (A) in the presence or absence of Oxy229 or Gefitinib, an EGFR tyrosine kinase inhibitor for 0, 40, and 60 min incubation. The upper pictures indicate the cells upon 40 min incubation at 37ºC in the presence or absence of the compounds. White arrows indicate the internalized speckles in colocalization with preS1, NTCP and EGFR. Merged pattern (right pictures) as well as nuclei (blue) are also shown. Scale bar, 10 μm. Subcellular localizations of preS1, NTCP and EGFR were observed at 0, 40 and 60 min after 37ºC incubation. Number of cells having internalized speckles for preS1, NTCP and EGFR was counted; percentages of those over the total counted cell numbers are shown as graphs. Statistical significance was determined by One-way ANOVA (**, P<0.01; *, P<0.05; N.S., not significant).

### Oxy229 attenuates the internalization of the preS1/NTCP complex, with remaining the intact EGFR endocytosis

We also detected the internalization of NTCP and EGFR as well as preS1 in the same assay, at 0, 40 and 60 min after starting 37ºC culture. Fig. 3B shows representative images at 40 min. As shown in Fig. 5A, NTCP and EGFR distributed at intracellular speckles in addition to cell surface, in colocalization with preS1 in DMSO-treated control cells (Fig. 3B, upper panel a-e). Although cell population showing such intracellular speckles for NTCP and EGFR was only 10-20% before 37ºC culture, it increased to 40-60% at 40 and 60 min after 37ºC incubation for DMSO-treated control (Fig 3B, lower graph, lanes 10, 13, 16, 19, 22, 25). As a positive control, in Gefitinib-treated cells, NTCP, EGFR and preS1 predominantly remained on cell surface even after EGF-stimulation, as reported (6) (Fig. 3B, upper panels k-o), with quantified internalized speckles significantly reduced at both 40 and 60 min (Fig. 3B, lower graph, lanes 6, 9, 15, 18 24, 27). In contrast, EGFR internalization was observed upon Oxy229 treatment, with no significant effect on quantified EGFR internalization at 40 and 60 min (Fig. 3B panel h and B, lanes 23, 26). However, intracellular speckles for preS1 and NTCP were clearly reduced by Oxy229 treatment, which was more drastic and significant than by Gefitinib (Fig. 3B panels f, g and B, lanes 5, 8, 14, 17). These data suggest that Oxy229 attenuated the internalization of the preS1/NTCP complex, with unaffected EGFR endocytosis, likely by dissociation of the preS1/NTCP complex from the relocating EGFR.

### Effect of Oxy229 on the original functions of NTCP and EGFR

We examined the effect of Oxy229 on the original functions of EGFR and NTCP. EGFR is implicated in variety of physiological functions such as cell proliferation and differentiation through activation of its downstream signaling (18). To assess the EGFR downstream signaling including Ras/MAPK and PI3K/Akt cascades, the cells were stimulated with EGF in the presence or absence of the compounds for 5 and 20 min, which were then analyzed to detect the phosphorylated form as well as the total protein of EGFR and its downstream Akt and ERK by immunoblot (Fig. 4A). In DMSO-treated control cells, EGF stimulation induced the phosphorylation of EGFR (Y1068), Akt (S473), and ERK (T202/Y204) (Fig. 4A, lane 2), and these phosphorylations were almost completely blocked upon Gefitinib treatment without affecting total EGFR, Akt, and ERK protein levels (Fig. 4A, lane 5), as expected (7). In contrast, Oxy229 and Oxy137 did not affect phosphorylations of EGFR, Akt, and ERK (Fig. 4A, lanes 3 and 4).

**Fig. 4.**
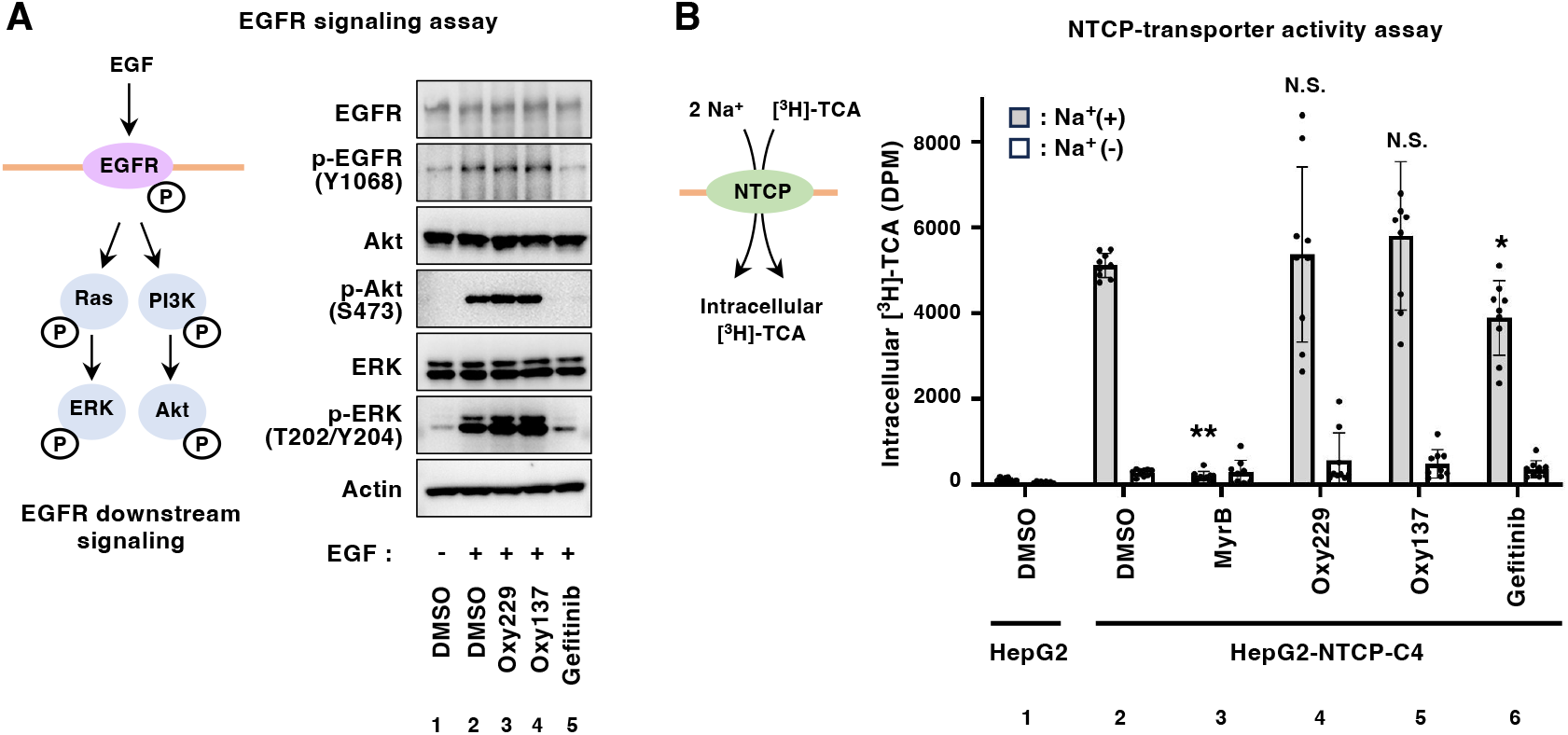
Effect of Oxy229 on the original functions of EGFR and NTCP. **(A)** Left, EGFR downstream signaling pathways. Right, HepG2-hNTCP-C4 cells were treated with the indicated compounds upon (+) or without (-) stimulation with 100 ng/ml EGF. Protein expressions of EGFR and its downstream Akt and ERK, and their phosphorylated forms [EGFR (Y1068), Akt (S473), and ERK (T202/Y204)] as well as actin as an internal control were detected by immunoblot. **(B)** Bile acid uptake activity of NTCP. HepG2-NTCP or HepG2 cells were incubated with [^3^H]-taurocholic acid (TCA) in the presence or absence of sodium along with the indicated compounds (0.9% DMSO, 100 nM MyrB, 90 μM Oxy229, 90 μM Oxy137, 30 μM gefitinib) for 15 min. After washing out free [^3^H]-TCA, intracellular radioactivity was counted with a liquid scintillation counter. Gray and white bars indicate the radioactivity in the presence and absence of sodium, respectively. Statistical significance was determined by One-way ANOVA (**, P<0.01; *, P<0.05; N.S., not significant).

NTCP is originally involved in the uptake of bile acids into the cells depending on sodium ion concentration gradient (16). We then examined the effect of the compounds on NTCP-mediated bile acid uptake by incubating the cells with [^3^H]-taurocholic acid as a substrate in the presence or absence of sodium upon treatment with or without compounds for 15 min and detecting the intracellular radioactivity (Fig. 4B). Upon DMSO treatment, the intracellular [^3^H]-taurocholic acid was detected in the presence, but not in the absence, of sodium in HepG2-hNTCP-C4 cells, indicating the sodium-dependent bile acid uptake, which was not seen in HepG2 cells that does not express endogenous NTCP (Fig. 4B, lanes 1 and 2). This intracellular uptake of [^3^H]-taurocholic acid was totally blocked by MyrB treatment, as reported (3, 19), but not by Gefitinib (Fig. 4B, lanes 3 and 6). In this assay, Oxy229 and Oxy137 did not significantly affect the uptake levels of [^3^H]-taurocholic acid (Fig. 4B, lanes 4 and 5) at the concentration ranges of Oxy229 showing anti-HBV activity (Fig. 2G). Thus, Oxy229 inhibited HBV infection without affecting the original function of EGFR and NTCP.

### An Oxy229 derivative, dually inhibiting both NTCP-EGFR interaction and NTCP multimerization, exhibits enhanced anti-HBV activity

To identify more potent anti-HBV compounds, we synthesized a small group of Oxy229 derivatives and examined their activities in infection assay using primary human hepatocytes. As structural backbone, the cholesterol and cholestanol units were employed, the latter lacking the C5-C6 double bond of cholesterol (Fig. 5A). We synthesized five derivatives by changing substituents in the sidechain attached at C-17 (R1 and R2), as shown in Fig. 5A. Each compound was treated at serially-diluted six concentrations (0.64, 1.6, 4, 10, 25, and 62.5 μM) to evaluate anti-HBV activity in the same HBV infection assay as shown in Fig. 3. In contrast to the parental Oxy229, exhibiting an IC_50_ of 13.0 μM in this assay, based on the reduction level of HBs antigen, Oxy234 and Oxy257, which R2 groups were chemically changed from biphenyl in Oxy229 to 3-pyridyl and *p*-(2-pheny1)phenyl, respectively, decreased the activity to inhibit HBV infection (IC_50_=37.1 and 47.8 μM, respectively) (Fig. 5A and B). In addition, Oxy284, possessing an acetylated hydroxyl groups at C-20 (R1) and at C-3, completely lost the activity (Fig. 5A and B), suggesting that free hydroxyl groups at C-20 (R1) and at C-3 may be necessary for the anti-HBV activity. Interestingly, Oxy283, which possesses the cholestanol backbone, devoid of the C5-C6 double bond, but is otherwise structurally identical to Oxy229, showed a remarkably elevated anti-HBV activity with an IC_50_ of 3.23 μM (Fig. 5A and B). Replacement of the 3-pyridyl in the R2 group on the cholestanol backbone (Oxy244), reduced the activity, as was case with the cholesterol, backbone-based modification from *p*-(2pyridy1)phenyl (Oxy229) to 3-pyridyl (Oxy234) that reduced anti-HBV activity (Fig. 5A and B). Analysis of these straightforward structure-activity relationships pointed to Oxy283 as an Oxy229 derivatives with increased anti-HBV potency.

**Fig. 5.**
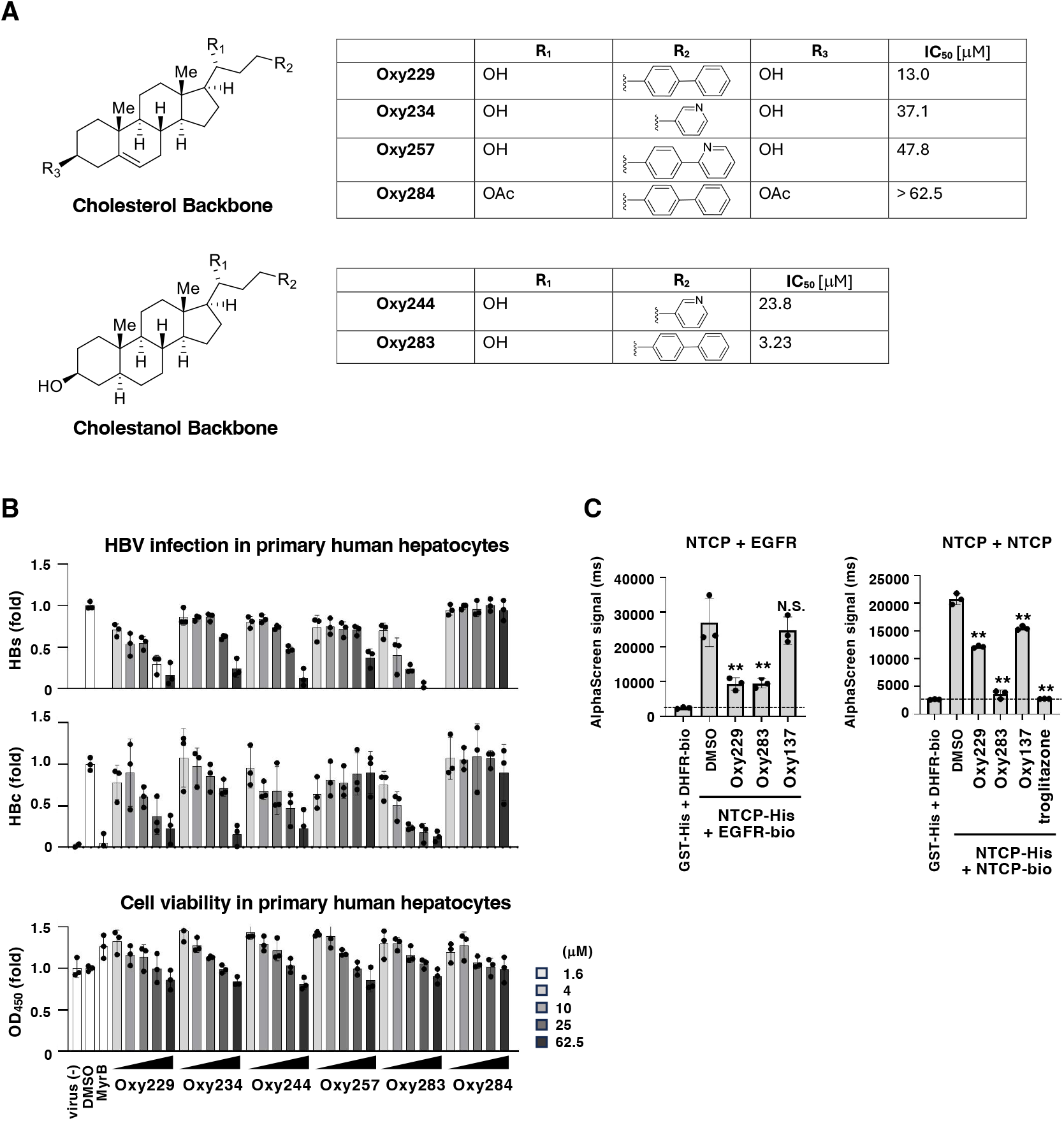
Anti HBV activity of Oxy229 derivatives. **(A)** Chemical structure of Oxy229 and its derivatives, based on their backbone structure, cholesterol or cholestanol, are shown on the left. Shown in the tables on the right, are compound name, substitution patterns in Rl and R2, IC_50_s for the inhibition of HBV infection. **(B)** HBV infection assay was performed using primary human hepatocytes upon treatment with or without the indicated compounds at varying concentrations (1.6, 4, 10, 25, and 62.5 μM). HBV infection was evaluated by measuring HBs antigen in the cultured supernatant (upper) and HBc antigen in the cells (middle). Cell viability was also measured by WST assay (lower). IC_50_s of these compounds are summarized in (A). **(C)** AlphaScreen assays for examining the interaction between His-tagged NTCP and biotinylated EGFR (left) and with His-tagged NTCP and biotinylated NTCP (right) in the presence of the indicated compound (DMSO, Oxy185, Oxy229, Oxy283, Oxy137, and troglitazone). The leftmost lanes show the co-incubation with GST-His and DHFR-bio to show the background signals (dotted lines). Statistical significance was determined by One-way ANOVA (**, P<0.01; *, P<0.05; N.S., not significant).

To get an insight into why Oxy283 had higher anti-HBV activity, we examined the effect of the compounds on the NTCP-mediated molecular interactions in the AlphaScreen assay. As shown in Fig. 5C, Oxy283 reduced the AlphaScreen signal produced by NTCP-EGFR binding to the similar level shown by treatment with Oxy229, in contrast to no significant effect of Oxy137 (Fig. 5C, left), in agreement with the similar inhibitory potential of Oxy283 and Oxy229 to NTCP-EGFR interaction. We also established the AlphaScreen assay evaluating the NTCP-NTCP interaction, another mechanism for facilitating HBV/HDV internalization (9) (Fig. 5C, right). In this assay, whereas Oxy229 only slightly affected the NTCP-NTCP interaction (to 55% of the contro1), Oxy283 more remarkably reduced NTCP multimerization (to 17%), as the level by treatment with troglitazone (to 13%), a reported NTCP multimerization inhibitor used as a positive control (9) (Fig. 5C, right). These data are consistent with that Oxy283 acquires the dual inhibitory activity to both NTCP-EGFR interaction and NTCP multimerization, thereby potentiating its anti-HBV activity compared to Oxy229. The implication of the dual inhibitory activity based on the chemical structure is discussed in the Discussion section below.

## Discussion

We have previously reported that EGFR supports the viral receptor function of NTCP as a cofactor through direct molecular interaction (7). EGFR directly interacts with NTCP either in the presence or absence of HBV/HDV, and its phosphorylation upon ligand stimulation triggers endocytosis together with NTCP and HBV/HDV. Since Oxy229 reduced the binding between recombinant EGFR and NTCP in vitro, it is likely to inhibit the molecular interaction by its direct binding to either NTCP or EGFR. From the results showing that Oxy229 inhibited HBV/HDV infection, whereas Oxy137, having no effect on NTCP-EGFR binding, had negligible antiviral effect, NTCP-EGFR interaction is suggested to play a relevant role in HBV/HDV infection.

Oxysterols are hydroxylated metabolites of cholesterol endogenously synthesized mostly by enzymes, such as cytochrome P450s. These naturally occurring oxysterols have variety of biological functions in lipid metabolism, signal transduction, and immune response, albeit with relatively weak activities, often paired with metabolic instability (20-22). We have developed and explored a series of unique semi-synthetic oxysterol derivatives, mainly through chemical modification of the sterol side chain, that resulted in improved drug-like properties in terms of potency, target selectivity, metabolic stability, safety, and oral bioavailability, with potential application in orthopedic medicine, chronic inflammation, fibrosis, cancer and infectious diseases such as COVID-19 (23-25). In this context, we previously reported that Oxy185, a cholestanol derivative, inhibited HBV entry without affecting the NTCP-EGFR interaction but by impeding the NTCP multimerization (10). Conceivably, the difference in chemical backbone between of Oxy185 and Oxy229 may be connected to their different modes of action with regard to the anti-HBV effects. Interestingly, Oxy283, identical to Oxy229 expect for its cholestanol backbone, displays greater anti-HBV potency compared to Oxy229 in primary human hepatocytes. We speculate that the increased potency of Oxy283 is resulted from dual inhibitory effects on both NTCP-EGFR interaction and NTCP multimerization, the two mechanisms of the viral receptor complex for supporting efficient HBV/HDV internalization. More in-depth derivative analysis in the future may identify more potent anti-HBV/HDV agents through the efficient blocking of this viral internalization machinery.

EGFR activates the downstream signaling including Ras-MAPK and PI3K-Akt pathways and are implicated in a variety of essential physiological events including cell proliferation and differentiation (18). NTCP is a hepatic transporter functioning for the bile acid uptake from the blood to hepatocytes and its genetic deficiency in mice exhibits phenotypes including the elevated serum bile acid levels, body weight loss, abnormal gallbladder morphology and aggravated osteoporosis (26-28). Initial clinical indications are that bulevirtide, when treated to HBV/HDV-coinfected patients, causes elevated bile acid levels but no severe adverse effects (29, 30). Nevertheless, it would be desirable to develop antiviral agents with limited effects on the original function of NTCP and EGFR. In this regard, our study demonstrates that Oxy229 selectively inhibits HBV/HDV entry without affecting the EGFR-mediated signaling and NTCP-mediated bile acid uptake (Fig. 6), making difference of Oxy229 from bluevirtide and Gefitinib to provide alternative approach for the development of a new type of anti-HBV/HDV agent.

**Fig. 6.**
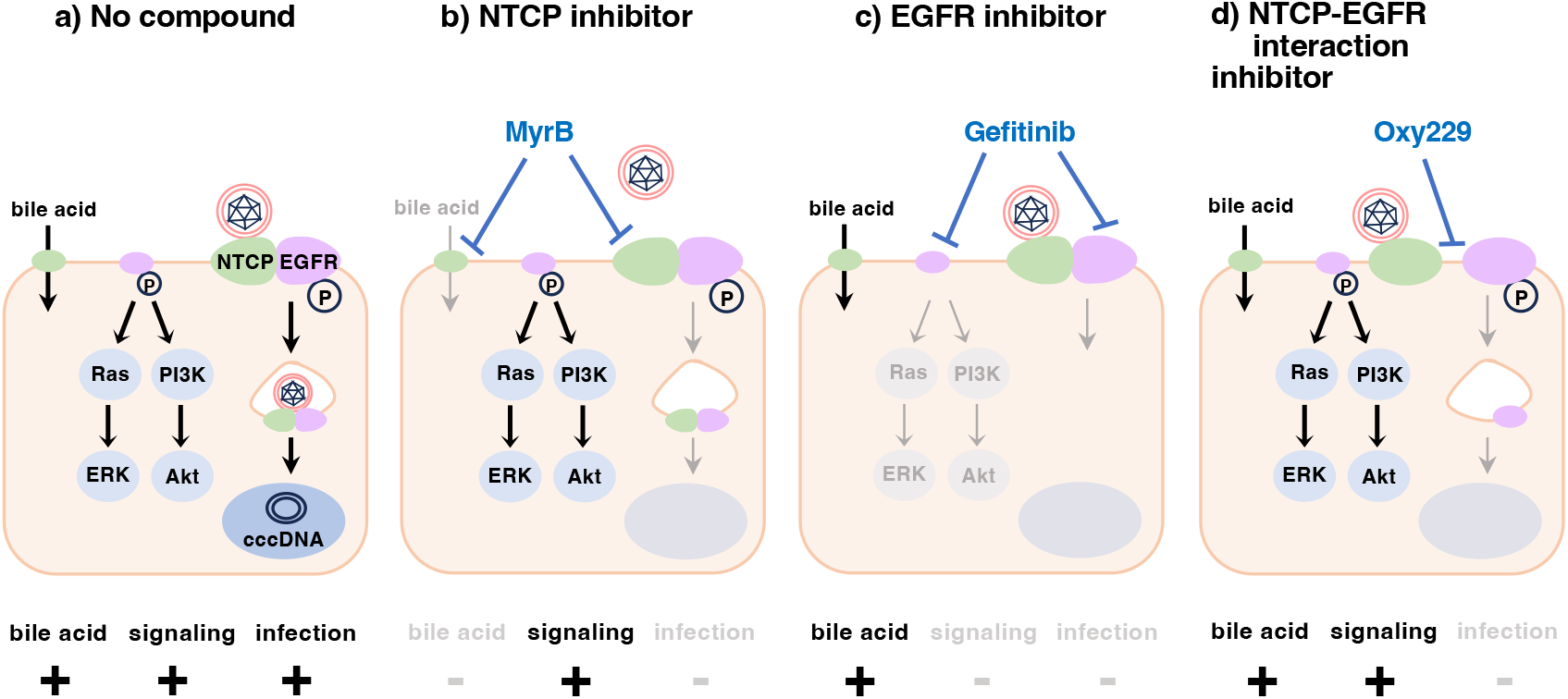
Schematic model of the mechanism for inhibiting HBV infection. **(A)** Without inhibitors, HBV enters into host cells in complex with NTCP and EGFR. NTCP and EGFR also function for the uptake of bile acid and activating the downstream signaling such as PI3K-Akt and Ras-MAPK pathways. **(b)** Myrcludex B inhibits HBV infection by blocking the preS1-NTCP interaction, and also inhibits the NTCP-mediated bile acid uptake. **(c)** EGFR antagonists such as Gefitinib inhibit the internalization of the HBV-NTCP-EGFR complex by blocking the phosphorylation of EGFR. Gefitinib also blocks the EGFR downstream signaling. **(d)** Oxy229 inhibited the interaction between NTCP and EGFR to inhibit HBV infection. Oxy229 does not affect the EGFR downstream signaling and NTCP-mediated bile acid uptake, thus limiting the adverse effects caused by inhibition of EGFR and NTCP physiological functions.

Thus, in this study, we identified Oxy229 and Oxy283, new compounds that disrupt the NTCP-EGFR interaction, as a novel mechanism inhibit HBV/HDV infection. In future studies, we plan to optimize antiviral activities and examine the pharmacokinetic properties of these oxysterol analogs, and the in vivo infection assays. However, this study at least shows that targeting the NTCP-EGFR-involved viral receptor complex offers a new approach for the development of antiviral agents.

## Materials and Methods

AlphaScreen assay was performed as previously described (11). HBV inoculum (genotype D) was prepared from the culture supernatant of Hep38.7-Tet cells as previously (17). The preS1 internalization assay was performed as previously described (6).

Detailed experimental procedures are described in SI Appendix, Supplementary Materials and Methods.

## Acknowledgements

The plasmids for expressing HDV and VSV pseudovirus were kindly provided by Dr. John Taylor at Fox Chase Cancer Center and Dr. Francois-Loic Cosset at University of Lyon. This work was supported by Japan Agency for Medical Research and Development (AMED) under grand numbers JP25Yk0310525 (K.W.), JP25Yk0310533 (K.W.), JP25Yk0319547 (K.W.), JP25ak0101271 (K.W.); JSPS/MEXT under KAKENHI grant numbers JP24K02290 (K.W.), JP25K21782 (K.W.), JP23K27415 (K.W.), JP25K21767 (K.W.), JP25K02691 (K.W.); JST MIRAI program grant number JPMJMI22G1 (K.W.); Takeda Science Foundation (K.W.)

## Notes

### Competing Interest Statement

F.S., F.W. and F.P are employees of Max BioPharma, Inc. R.M. is an employee of Cell Free Sciences Co., Ltd.

