## Supplementary material for "Disruption of the NTCP-EGFR receptor complex as a strategy for selective inhibition of hepatitis B and D virus entry": SI

**Contents:**

Supplementary Materials and Methods
Supplementary References
Supplementary Figure legends
Supplementary Figures

#### Supplementary Materials and Methods

##### AlphaScreen

Recombinant His-tagged NTCP (NTCP-His) and biotinylated EGFR (Bio-EGFR) proteins were synthesized in wheat germ cell free protein expression system (Cell Free Sciences) as previously described (1, 2). The primary screening was performed by co-incubating NTCP-His and Bio-EGFR in the buffer [100 mM Tris-HCl (pH 8.0), 150 mM NaCl, 0.01% NP40, 1 mg/ml BSA] with addition of streptavidin-conjugated donor beads and anti-His-conjugated acceptor beads in the presence of 100 μM compounds from the chemical library for 1 h, and AlphaScreen signal was detected with a microplate reader as described (1). The chemical library consists of 1,000 compounds including in-house small molecules and peptides. As the secondary assay to eliminate pseudo-positive compounds, biotinylated-His tag peptide (Bio-His, PerkinElmer) was incubated with the streptavidin-coated donor beads and anti-His-conjugated acceptor beads in the same buffer in the presence of 100 μM compounds.

##### Cell culture

HepG2 and HepG2-hNTCP-C4 cells (3) were cultured with DMEM/F12 Glutamax supplement (Gibco) supplemented with 10 mM HEPES (Nacalai), 100 units/ml penicillin, 100 μg/ml streptomycin, 10% FBS, and 5 μg/ml insulin in the presence (HepG2-hNTCP-C4 cells) or absence (HepG2 cells) of 400 μg/ml G418 (Nacalai). Hep38.7-Tet cells (4) were cultured in the above medium supplemented with 0.4 μg/ml tetracycline during passages. Huh-7 cells were cultured in D-MEM (High Glucose, Wako) supplemented with 10 mM HEPES, 100 units/ml penicillin, 100 μg/ml streptomycin, 10% FBS, 100 μM non-essential amino acids, and 1 mM sodium pyruvate. Primary human hepatocytes (PHH) (PhoenixBio Co, Ltd) were maintained in the medium purchased from PhoenixBio, Inc. All the cells were cultured at 37°C, 5% $CO_2$. In the assay using EGF stimulation, the cells were precultured in the medium without FBS and insulin for 24 h and were stimulated with 10 or 100 ng/ml EGF in the medium without FBS.

##### Reagents and compounds

EGF was purchased from PeproTech. Mycludex B and Entecavir are obtained from Selleck and Santa Cruz respectively. Gefitinib and Dimethyl sulfoxide (DMSO) were purchased from Sigma-Aldrich. TAMRA-labeled preS1 peptide (preS1-TAMRA) was synthesized by Scrum. The chemical library consists of 1,000 compounds including in house small molecules and peptides. Oxysterols were synthesized according to well-established methods and procedures we previously described (5-7).

##### HBV infection assay

HBV inoculum (genotype D) was prepared from the culture supernatant of Hep38.7-Tet cells as previously described (8). HepG2, HepG2-hNTCP-C4 cells or PHH were inoculated with HBV at 250-3,000 genome equivalent (GEq)/cell in the presence of 4% PEG 8000 for 16 h. After washing out free virus and compounds, the cells were cultured for an additional 12 days in the absence of compounds. HBV infection was evaluated by detecting either HBs antigen in the culture supernatant, HBc antigen and HBV DNA in the cells by ELISA, immunofluorescence, and real-time RT-PCR, respectively.

**ELISA**

HBs antigen was quantified by ELISA using plates immobilized with anti-HBs antibody (Abcam) in 0.2% BSA, 0.02% $NaN_3$, 1 x PBS at 4°C. Culture supernatants were added to the plates for 2 h, followed by incubation with horse radish peroxidase-labeled rabbit anti-HBs antibody (LSBio) for 2 h and visualization with the substrate solution (Sumitomo Bakelite) to measure the absorbance at 450 nm by multiple reader (Tecan), as described previously (8).

**Immunofluorescence for detecting HBV infection**

HBc antigen was detected by indirect immunofluorescence analysis for evaluating HBV infection. After fixation with 4% paraformaldehyde, the cells were permeabilized with 0.3% Triton X-100 and then were treated with an anti-HBc antibody (Beacle) as a primary antibody, followed by treatment with Alexa Fluor 555-conjugated Goat anti-rabbit antibody (Invitrogen) together with 0.02% 4'6-diamidino-2-phenylindole (DAPI) for visualizing HBc antigen and the nuclei. These cells were observed by a fluorescence microscopy (BZ-X800, Keyence) (8).

**Real-time PCR**

HBV DNA was extracted by QIAamp DNA Mini Kit (QIAGEN) according to the manufacturer's protocol. Real-time PCR for the quantification of HBV DNA was performed using primer-probe set: 5'-AAGGTAGGAGCTGGAGCATTCG-3' and 5'-AGGCGGATTTGCTGGCAAAG-3' as primer sets and 5'-[FAM]AGCCCTCAGGCTCAGGGCATAC[TAMRA]-3' as a probe. The reaction condition was 95°C for 10 min, 50 cycle of 95°C for 15 sec, and 60°C 1 min for 50 cycles (8).

**HBV replication assay**

HBV replication activity was evaluated by Hep38.7-Tet cells, which can induce HBV replication from an HBV transgene integrated in the host chromosome by depletion of tetracycline from the medium (4). The cells were treated with or without the compounds in the absence of tetracycline for six days, with changing the medium to that containing the compounds but not tetracycline every three days. DNA in the supernatant was recovered and HBV DNA was quantified by real-time RT-PCR (8).

**Cell viability**

Cell viability was evaluated by WST assay using Cell counting kit-8 (DOJINDO), according to the manufacturer's instruction, as described (9).

**PreS1 binding assay**

To evaluate preS1 mediated cell attachment, HepG2-hNTCP-C4 cells were treated with 40 nM C-terminally TARMA-conjugated and N-terminally myristoylated peptide spanning 2-48 aa of the preS1 (preS1-TAMRA) at 4°C for 30 min to allow cell surface attachment but not intracellular internalization, as described (8). After fixation with 4% paraformaldehyde and staining with DAPI, the cells were observed with a fluorescence microscopy (BZ-x800, Keyence).

**Internalization assay**

HepG2-hNTCP-C4 cells and HepG2 cells stably overexpressing NTCP fused with oxy StayGold, were treated with 40 nM preS1-TAMRA at 4°C for 30 min to allow cell attachment, and were then transferred to incubate at 37°C by treatment with 10 ng/ml EGF to stimulate internalization, as described (10). The cells were fixed, permeabilized, and treated with anti-EGFR antibody (Cell signaling) as a primary antibody, followed by treatment with Alexa Fluor 647-conjugated Goat anti-rabbit antibody (Invitrogen) together with DAPI to observe with a confocal microscopy (LSM 900, ZEISS). For quantification of the percentage of cells showing the internalization of preS1, NTCP and EGFR, the cell exhibiting five or more intracellular speckles was defined as an internalized cell and the number of internalized cells was counted among >100 cells for each sample to calculate the percentage of internalized cells, as the method described previously (10).

**Immunoblot analysis**

Activation of the EGFR downstream signaling was examined by immunoblotting. Cell lysates were fractionated by SDS-page, transferred to a PVDF membrane (Immobilion-P, Millipore), and subjected to immunoblot with anti-EGFR (Cell Signaling), anti-phospho EGFR (Y1068) (Cell Signaling), anti-Akt (CELL Signaling), anti-phospho Akt (S473) (Cell Signaling), anti-ERK (Cell Signaling), anti-phospho ERK (T202/Y204) (Cell Signaling) and anti-actin (sigma-Aldrich) antibodies as primary antibodies. These immunocomplex were visualized by incubating with HRP-conjugated donkey anti-rabbit IgG antibody (Cell Signaling) and the chemiluminescent substrate (Thermo Science), and observed with Chemiluminescence imaging system (WSE-6370, ATTO).

**NTCP transporter assay**

NTCP-mediated bile acid uptake activity was measured by preincubating HepG2 or HepG2-hNTCP-C4 cells with the compounds for 1 h and then incubating them with [$^3$H]-taurocholic acids (TCA) in the buffer (4.8 mM KCl, 1.2 mM $KH_2PO_4$, 1.2mM $MgCl_2$, 1.3 mM $CaCl_2$, 2.6 mM D-glucose, 25 mM HEPES, 10 μM TCA, pH 7.4) with or without 125 mM sodium in the presence or absence of the compounds, to allow [$^3$H]-TCA uptake into the cells at 37°C for 10 minutes. After washing the free [$^3$H]-TCA, the cells were lysed by 0.05% SDS and the intracellular radioactivity was measured with a liquid scintillation counter (Tri-Carb 2810TR, TR-LSC), as described (8).

**HDV infection assay**

HDV inoculum was prepared by recovering the culture supernatant of Huh-7 cells transfected with the plasmid encoding HDV (kindly provide by Dr. John Taylor at Fox Chase Cancer Center) and that encoding HBs antigen, as described (8). HepG2-hNTCP-C4 cells were inoculated with HDV at 125 GEq/cell with 5% PEG8000 for 16 h, washed, and cultured for five days. Intracellular HDV RNA was detected by real time RT-PCR to evaluate HDV infection (8).

**Vesicular stomatitis virus (VSV) entry assay**

VSV entry to cells was evaluated with a retrovirus-based VSV pseudovirus carrying VSV G protein as an envelope protein on the particles. VSV pseudovirus was prepared from culture supernatant of 293T cells transfected with the expression plasmids encoding MLV Gag-pol, luciferase, and VSV G (kindly provided by Dr Francois-Loic Cosset at University of Lyon), as described (11). VSV-mediated entry was evaluated by treatment of Huh-7 cells with the VSV pseudovirus together with or without the compounds for 1 h and incubating the cells for 72 h to measure the luciferase activity (11).

**Real time RT-PCR**

HDV RNA was extracted by RNeasy Mini Kit (Qiagen) according to the manufacturer's protocol and was detected by High-capacity cDNA RNase Transcription Kit (fisher scientific) with RNase Inhibitor (Fisher Scientific). PCR for the quantification of HDV RNA was performed by using a primer-probe set: 5'-GGACCCCTTCAGCGAACA-3' and 5'-CCTAGCATCTCCTCCTATCGCTAT-3' as primers and 5'-AGGCGCTTCGAGCGGTAGGAGTAAGA-3' as a probe. PCR reaction was performed with 40 cycles of 95°C for 5sec, and 60°C for 34 sec for 40 cycles. (8) .

**Statistics**

Statistical significance was determined by One-way ANOVA (**, $p<0.01$; *$P<0.05$; N.S., not significant) using GraphPad Prism.

#### Supplementary Figure legends

**Fig. S1.** Oxy229 showed anti-HBV activity in primary human hepatocytes.
**(A-C)** HBV infection assays were performed as shown in Fig. 2B with primary human hepatocytes in the presence or absence of the indicated compounds [0.9% DMSO, 100 nM MyrB, 120 μM Oxy229, 500 nM ETV, 50 μM Gefitinib]. HBV infection was determined by detecting HBs antigens in the cultured supernatant (A) and HBc antigen in the cells (C). Red and blue signals in (C) indicate HBc antigen and nuclei, respectively. Scale bar, 100 μm. Cell viability was also measured by WST assay (B).

**Fig. S2.** Oxy229 did not affect HBV attachment and replication.
**(A)** Schematic representation of the HBV lifecycle. HBV attaches to the cells through the NTCP binding (attachment), followed by internalization inside the cells (internalization) that leads to the translocation into the nucleus and eventually to form covalently closed circular DNA (cccDNA). HBV replication is initiated to transcribe from cccDNA into HBV RNAs and produces viral particles (replication). Activities of replication, attachment, and internalization were evaluated in Fig. S2B, Fig. S2C, and Fig. 3A, respectively. **(B)** HBV replication assay. Hep38.7-Tet cells, which induces HBV replication upon depletion of tetracycline, were treated with compounds in the absence of tetracycline for six days with changing medium every three days, and HBV DNA in the cultured supernatant was quantified by real-time RT-PCR. **(C)** HBV attachment was evaluated with TAMRA-labeled N-terminally myristoylated preS1 peptide consisting of 2-48 amino acid region (preS1-TAMRA). HepG2 or HepG2-hNTCP-C4 cells were treated with preS1-TAMRA in the presence or absence of compounds at 4°C for 30 min. After washing and fixation, the cells were detected for preS1-TAMRA (red) as well as nuclei (blue) with a fluorescence microscopy. Fluorescence areas were quantified using an image analyzer (BZ-X800, KEYENCE) and are shown as a graph on the right.

#### Fig. S1

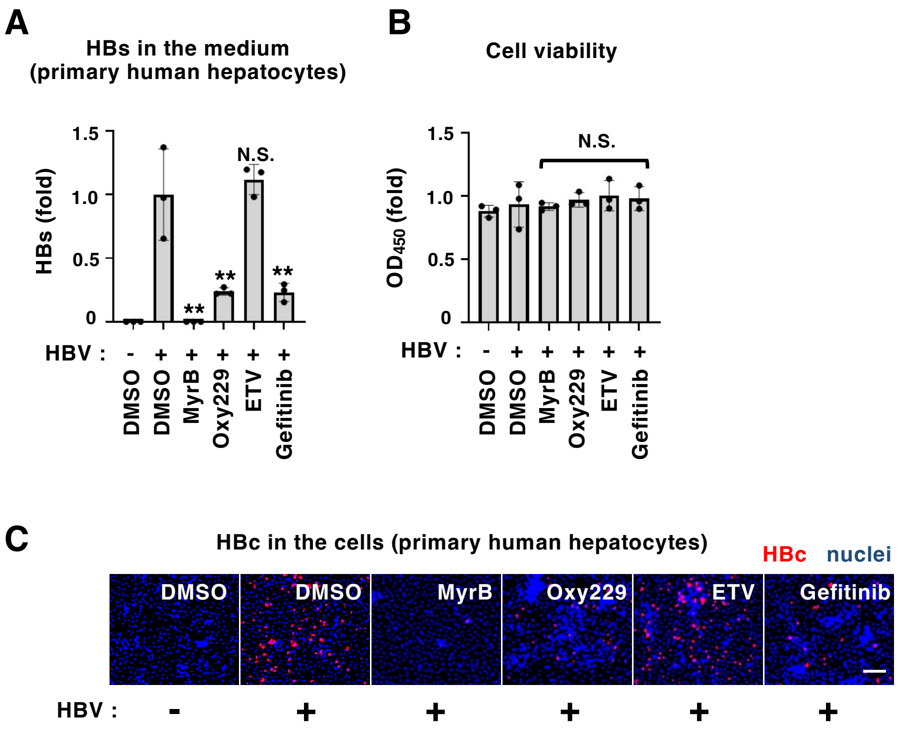
A
HBs in the medium
(primary human hepatocytes)
HBs (fold)
1.5
1.0
0.5
0
N.S.
**
**
**
HBV : - + + + + +
DMSO
DMSO
MyrB
Oxy229
ETV
Gefitinib
B
Cell viability
OD450 (fold)
1.5
1.0
0.5
0
N.S.
HBV : - + + + + +
DMSO
DMSO
MyrB
Oxy229
ETV
Gefitinib
C
HBc in the cells (primary human hepatocytes)
HBc
nuclei
DMSO
DMSO
MyrB
Oxy229
ETV
Gefitinib
HBV : - + + + + +

### Fig. S2

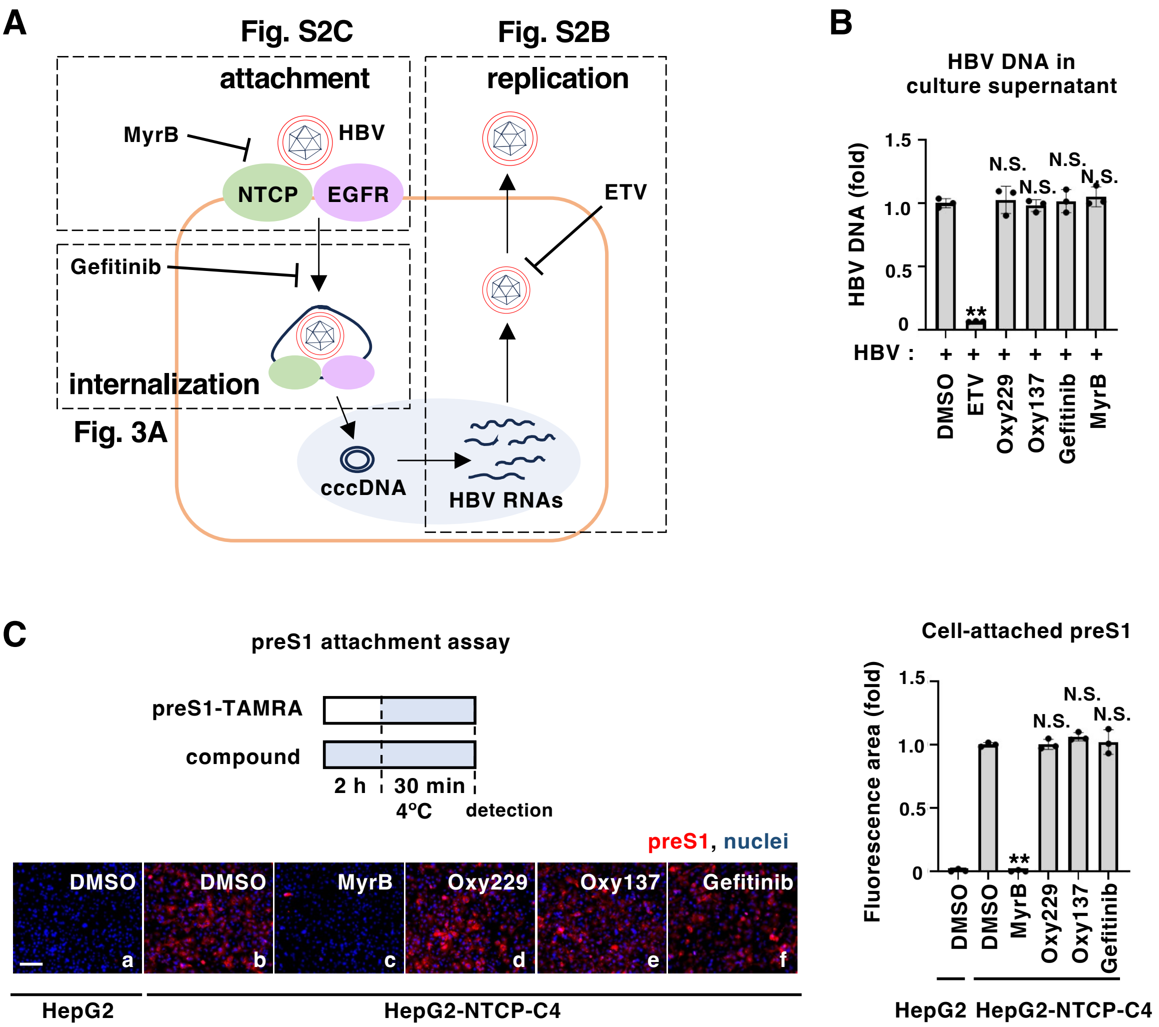
A
Fig. S2C
attachment
MyrB
HBV
NTCP
EGFR
Gefitinib
internalization
Fig. 3A
Fig. S2B
replication
ETV
cccDNA
HBV RNAs
B
HBV DNA in culture supernatant
HBV DNA (fold)
1.5
1.0
0.5
0
N.S.
**
HBV :
+
DMSO
ETV
Oxy229
Oxy137
Gefitinib
MyrB
C
preS1 attachment assay
preS1-TAMRA
compound
2 h
30 min
4°C
detection
preS1, nuclei
DMSO
DMSO
MyrB
Oxy229
Oxy137
Gefitinib
a
b
c
d
e
f
HepG2
HepG2-NTCP-C4
Cell-attached preS1
Fluorescence area (fold)
1.5
1.0
0.5
0
N.S.
**
DMSO
DMSO
MyrB
Oxy229
Oxy137
Gefitinib
HepG2
HepG2-NTCP-C4
